# CRISPR/Cas9-mediated genome editing reveals the involvement of a polyphenol oxidase in the shikonin-specific biosynthesis in *Lithospermum erythrorhizon*

**DOI:** 10.1101/2025.07.30.667576

**Authors:** Kohei Nakanishi, Yuki Takano, Kyoko Yamamoto, Mariko Yano, Koji Mito, Takuji Ichino, Kanade Tatsumi, Hao Li, Kazuaki Ohara, Ryosuke Munakata, Hideyuki Suzuki, Nozomu Sakurai, Daisuke Shibata, Keishi Osakabe, Bunta Watanabe, Takahiro Okada, Koichiro Shimomura, Kojiro Takanashi, Akifumi Sugiyama, Kazufumi Yazaki

**Affiliations:** Research Institute for Sustainable Humanosphere, Kyoto University, Gokasho, Uji, Kyoto 611-0011, Japan; Institute of Natural Medicine, Toyama University, 2630 Sugitani, Toyama, Toyama 930-0194, Japan; Laboratory of Medicinal Cell Biology, Kobe Pharmaceutical University, 4-19-1 Motoyamakita, Higashinada, Kobe, Hyogo 658-8558, Japan; Kazusa DNA Research Institute, Kisarazu, Chiba 292-0818, Japan; Graduate School of Technology, Industrial and Social Science, Tokushima University, 3-18-15 Kuramoto, Tokushima, Tokushima 770-8503, Japan; Chemistry Laboratory, The Jikei University School of Medicine, 8-3-1 Kokuryo, Chofu, Tokyo 182-8570, Japan; Division of Molecular Cell Biology, Department of Biomolecular Sciences, Faculty of Medicine, Saga University, Japan; Graduate School of Life Science, Toyo University, 1-1-1 Izumino, Itakura, Ora, Gunma 374-0193, Japan; Graduate School of Science and Technology, Shinshu University, 3-1-1 Asahi, Matsumoto, Nagano 390-8621, Japan

**Keywords:** shikonin biosynthesis, 1, 4-naphthoquinone derivatives, copper ion, polyphenol oxidase, *Lithospermum erythrorhizon*, red gromwell

## Abstract

Shikonin, a 1,4-naphthoquinone derivative produced by several Boraginaceae species, exhibits unique pharmacological properties and is used as a natural dye. The regulatory factors of shikonin production have been demonstrated using a cell culture system of *Lithospermum erythrorhizon*. Among these factors, copper is known to be the strongest enhancer of shikonin production. Although shikonin biosynthesis has been studied for over 40 years, the steps of naphthalene ring formation are still unknown, as is the reason for the effect of copper. In this study, we explored candidate genes associated with shikonin production using a PCR-select subtraction experiment. Polyphenol oxidase (PPO), a dicopper-dependent oxidoreductase, was highlighted because it showed synchronous expression with shikonin production. Transcriptome analysis of hairy roots and cultured cells of this plant revealed that, of the five *PPO* genes expressed in *L. erythrorhizon*, only *PPO1* showed a strong correlation with shikonin production. Next, we generated genome-edited hairy roots of *LePPO1* using CRISPR/Cas9-mediated mutagenesis to analyze its impact on shikonin derivative and other specialized metabolite production. The results showed that shikonin content was markedly reduced in all *LePPO1*-ge lines. Interestingly, the content of deoxyshikonofuran, a hydroquinone derivative and shunt product that branches after GHQ-3′′-OH in the shikonin biosynthetic pathway, remained unaffected in the *LePPO1*-ge lines. These findings suggest that LePPO1 participates in naphthalene ring formation and explain why a copper ion is crucial for shikonin biosynthesis.

## Introduction

Plant specialized metabolites are widely used in human life. They are used as functional foods, medicinal materials, cosmetics, and dyes for clothing. Shikonin derivatives, which are red 1,4-naphthoquinones, are produced only by a few species in the Boraginaceae family. Red gromwell (*Lithospermum erythrorhizon*) is one such species. These derivatives have been studied extensively in the field of pharmacology due to their various beneficial properties, including anti-inflammatory, antitumor, antibacterial, and granulation-promoting activities (Sankawa et al. 1977, Tanaka et al. 1986, Ozaki et al. 1998, Brigham et al. 1999, Guo et al. 2019). They have also been reported to inhibit SARS-CoV-2 main protease (Jin et al. 2020a, Jin et al. 2020b). Red gromwell is also known for its use as a natural dye, and its roots have been used to dye fabrics and Japanese kimonos (Yazaki 2017). Moreover, several recent studies have reported that plant-microbe interactions via shikonin derivatives promote the growth and production of shikonin in shikonin-producing species (Alonso et al. 2022, Rat et al. 2023). However, *L. erythrorhizon* is an endangered species due to climate change, overharvesting, and the spread of plant-pathogenic viruses (Izuishi et al. 2020, Ito et al. 2023). Fortunately, a cell culture system for this plant was established in the 1970s and 1980s, and the regulatory factors of shikonin production were clarified using cultured cells. Positive regulators of shikonin production include methyl jasmonate, copper ions (Cu²⁺), and low temperatures (<21 °C), while negative regulators include light irradiation (especially blue light), ammonium ions (NH₄⁺), and high temperatures (>28 °C) (Yazaki, 2017). However, the molecular mechanisms of these regulatory factors remain unclear. In particular, the reason for the requirement of copper ions has drawn significant attention from researchers due to its striking effect.

Since the genome assembly report on *L. erythrorhizon* (Auber et al. 2020, Fu et al. 2021, Tang 2021, Suttiyut et al. 2022, Li et al. 2022, Nakanishi et al. 2024, Okada and Watanabe 2024), the shikonin biosynthetic genes have been successively identified by a reverse genetic approach. Shikonin biosynthesis involves two key precursors: *p*-hydroxybenzoic acid (PHB), which is derived from the phenylpropanoid pathway, and geranyl diphosphate (GPP), which is derived from the mevalonate pathway (Ueoka et al. 2020). These precursors are coupled by *p*-hydroxybenzoate:geranyltransferase (PGT) (Yazaki et al. 2002) to form *m*-geranyl-*p*-hydroxybenzoic acid (GBA) (Fig. 1). Then, 3′′-hydroxygeranylhydroquinone (GHQ-3′′-OH) forms via the decarboxylation and hydroxylation of GHQ by GHQ 3′′-hydroxylase (GHQH) (Yamamoto et al. 2000a, Song et al. 2020). GHQ-3′′-OH then proceeds to the naphthalene ring formation step. This step is followed by side-chain modification to produce shikonin derivatives (Oshikiri et al. 2020, Song et al. 2021, Oshikiri et al. 2024). Although the final two steps of the shikonin biosynthetic pathway have been discovered, the naphthalene ring formation step remains unclear (Widhalm & Rhodes 2016).

**Figure 1.**
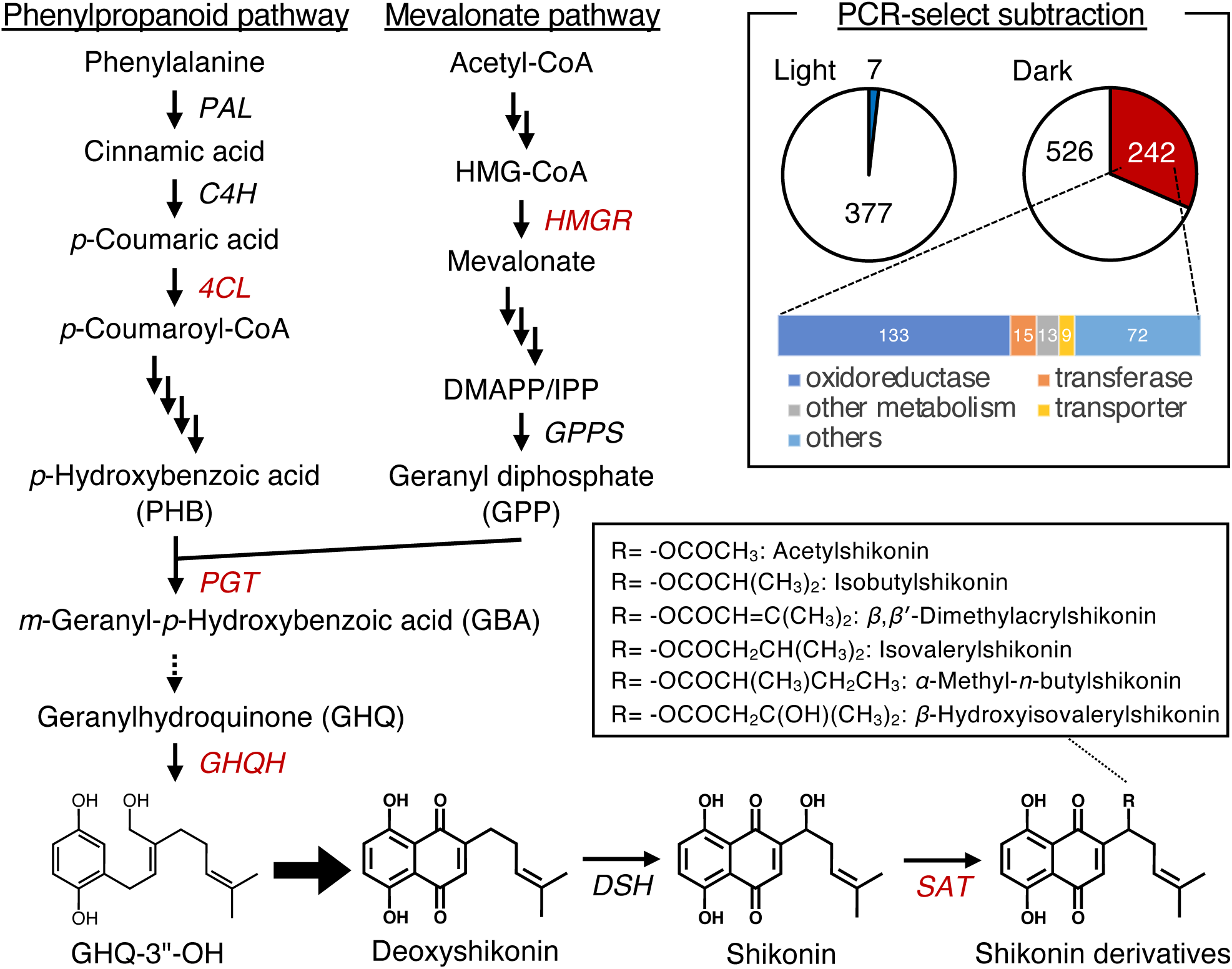
Overview of the shikonin biosynthetic pathway and biosynthetic genes induced under dark condition in *Lithospermum erythrorhizon* cultured cells. The shikonin biosynthetic genes highlighted as dark-induced genes in PCR-select subtraction are shown as red letters. Each arrows represents a single biosynthetic reaction, dashed arrow represents vailed reaction, and thick arrow is the target reaction steps in this study. Gene names and abbreviations are shown as follows: *PAL*, *phenylalanine ammonia lyase*; *C4H*, *cinnamic acid 4-hydroxylase*; *4CL*, *4-coumaroyl-CoA ligase*; *HMGR*, *3-hydroxy-3-methyl-glutaryl-CoA reductase*; *GPPS*, *geranyl diphosphate synthase*; *PGT*, *p-hydroxybenzoate:geranyltransferase*; *GHQH*, *geranylhydroquinone* 3′′*-hydroxylase*; *DSH*, *deoxyshikonin hydroxylase*; *SAT*, *shikonin O-acyltransferase*.

In this study, we used the PCR-select subtraction method to explore candidate genes involved in naphthalene ring formation in the shikonin biosynthetic pathway from dark-induced genes expressed in cultured *L. erythrorhizon* cells. Through expression profiling and phylogenetic analysis, we identified a gene that encodes polyphenol oxidase, a copper enzyme widely distributed from bacteria to plants. We took this gene, a strong candidate in shikonin production, and generated CRISPR/Cas9-mediated genome editing mutants to demonstrate its involvement in naphthoquinone formation, but not in hydroquinone-type shunt products. These findings advance our understanding of the regulatory mechanism of copper ions and the formation of the naphthoquinone skeleton in shikonin biosynthesis.

## Results

### *LePPO1* is highlighted as a dark-induced gene with shikonin biosynthetic genes

Shikonin production is induced by culturing in the M9 pigment production medium under dark conditions (Yazaki et al. 1999). To identify the genes involved in shikonin production, we cultured *L. erythrorhizon* (TOM strain) cells in an M9 medium under light (no shikonin production) or dark (shikonin production) conditions and performed PCR-select subtraction analysis. Among the 384 gene fragments found to be expressed in TOM cells cultured under illumination, seven were identified as light-induced genes (Fig. 1). In contrast, 768 gene fragments were cloned as transcripts in TOM cells in the dark. Of these, 242 were identified as dark-induced genes. These genes were grouped according to their function as follows: 133 oxidoreductases, 15 transferases, 13 genes related to other metabolisms, nine transporters, and 72 other genes (Fig. 1). The list of dark-induced genes included the following known genes involved in shikonin biosynthesis: *4-coumaroyl-CoA ligase* (*4CL*), *3-hydroxy-3-methylglutaryl-CoA reductase* (*HMGR*), *PGT*, *GHQH*, and *shikonin O-acyltransferase* (*SAT*). Notably, 125 of the 242 dark-induced genes showed induction rates of 100-fold or greater. Of those, 17 gene fragments were annotated as *PPO* genes encoding copper-dependent oxidoreductases (Fig. 1). These gene fragments were found to be derived from one gene. The cDNA was cloned from a cDNA pool prepared from cells cultured in an M9 medium in the dark and designated *LePPO1*. Because copper is essential for shikonin production, *LePPO1* was investigated as a primary candidate in further studies.

### The expression of *LePPO1* is associated with shikonin production in *L. erythrorhizon*

Polyphenol oxidase (PPO; EC 1.10.3.1) is widely distributed in the plant kingdom and is also found in bacteria and fungi (Mayer 2006). The representative reaction that it catalyzes is the hydroxylation and oxidation of monophenols and diphenols, which uses oxygen as a substrate (Yoruk and Marshall 2003, Tran et al. 2012). The browning of crops and apples is a well-known example of this reaction. Note that the PPO family includes the laccase family, which is relevant for lignin polymerization (Liang et al. 2006, Zhang et al. 2013). These laccases are grouped in a separate clade (Bai et al. 2023). Using LePPO1 as a query, we surveyed a gene database to determine if LePPO1 belongs to the typical PPO group (catechol oxidase or tyrosinase) or the laccase group. We found that LePPO1 belongs to the former group (Supplemental Fig. S1). Further analysis of the *L. erythrorhizon* transcriptome revealed the expression of four additional *PPO* genes. To analyze the expression profiles of these five *PPO* genes in cultured TOM cells and *L. erythrorhizon* hairy roots in M9 medium under light or dark conditions, we performed a detailed transcriptome analysis in comparison to the 13 previously identified shikonin biosynthetic genes. The analysis revealed that, of the five *LePPOs*, only *LePPO1* was notably expressed under dark conditions alongside other shikonin biosynthetic genes in both cultured cells and hairy roots (Fig. 2A). Shikonin derivatives are produced specifically in root tissues, particularly in the root epidermis (Yamamoto et al. 2000b). We evaluated the relative expression levels of *LePPO1* in various organs of intact *L. erythrorhizon* plants to clarify the organ specificity of *LePPO1* expression with shikonin production. The results showed that *LePPO1* is highly expressed only in organs where shikonin is produced; almost no expression was observed when the shikonin-containing peel was removed, even from the roots (Fig. 2B). These results suggest that *LePPO1* is coordinately expressed with shikonin biosynthetic activity.

**Figure 2.**
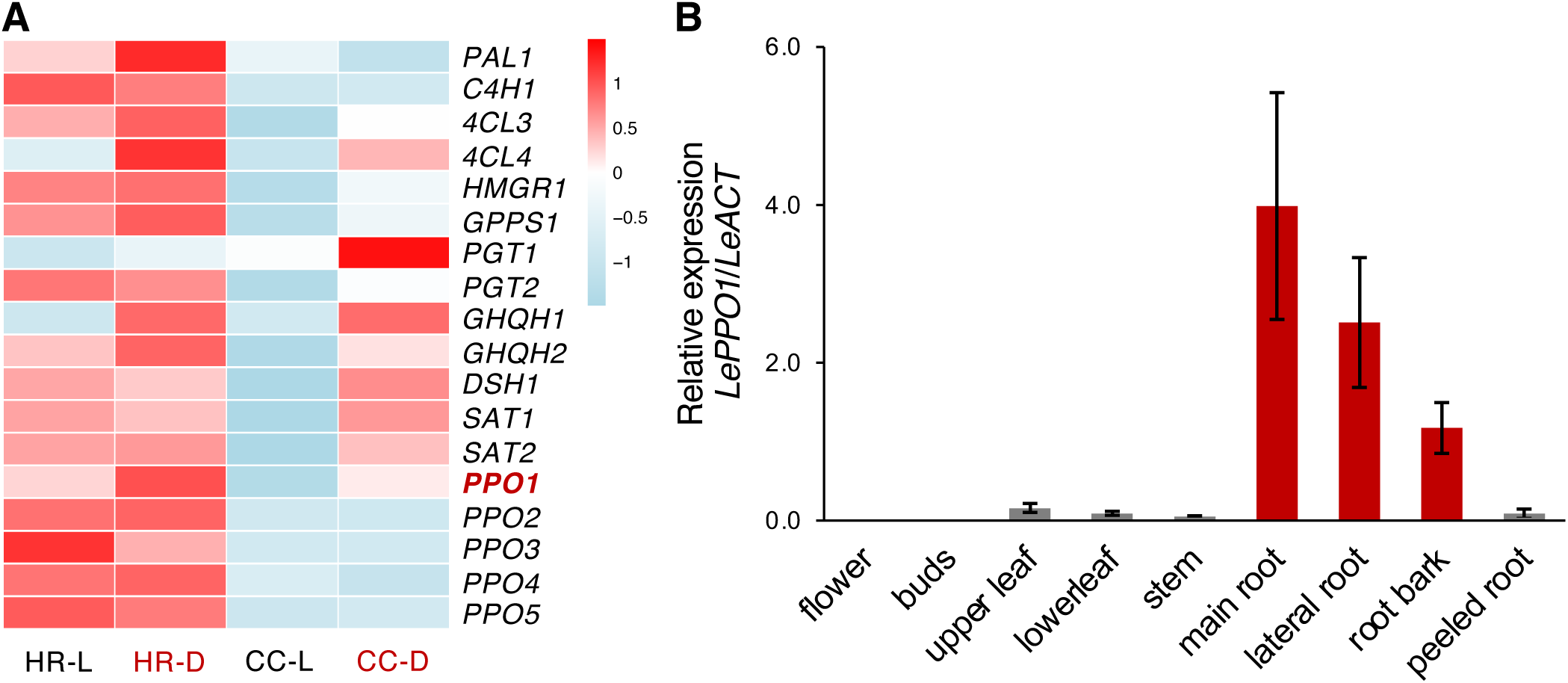
Expression profile of five *LePPO* genes in *L. erythrorhizon*. A: Expression profiles of five *LePPOs* and other known shikonin biosynthetic genes in hairy roots and cultured cells cultured under the light or dark conditions. Transcripts per million (TPM) were converted to log2 (TPM+1) and normalized by Z-scores. The abbreviations of samples are shown as follows: HR-L, hairy roots-light; HR-D, hairy roots-dark; CC-L, cultured cells-light; CC-D, cultured cells-dark. B: Relative expression levels of *LePPO1* in different organs of *L. erythrorhizon*. *LeACT* was used as an internal standard gene. Bars are represented as means ± SD of three technical replicates. Red bars indicate the tissues where shikonin production occur.

### LePPO1 having two conserved copper-binding domains is classified in a family-specific clade

PPO is a dicopper-dependent protein with two conserved regions that interact with copper ions, called CuA and CuB, which have three His residues, respectively (Tran et al. 2012). Figure 3 shows the multiple sequence alignment of LePPO1 with several representative plant PPOs (Liao et al. 2006, Yu et al. 2020, Wei et al. 2022, Sajjad et al. 2023). The figure depicts the presence of the CuA and CuB domains in LePPO1, which include three His residues: H197, H221, and H230 in the CuA domain, as well as H371, H375, and H404 in the CuB domain (Fig. 3A). These findings suggest that LePPO1 functions as a copper-dependent oxidase. AlphaFold 3 predicted the 3D structure of LePPO1 with two copper ions (Fig. 3B).

**Figure 3.**
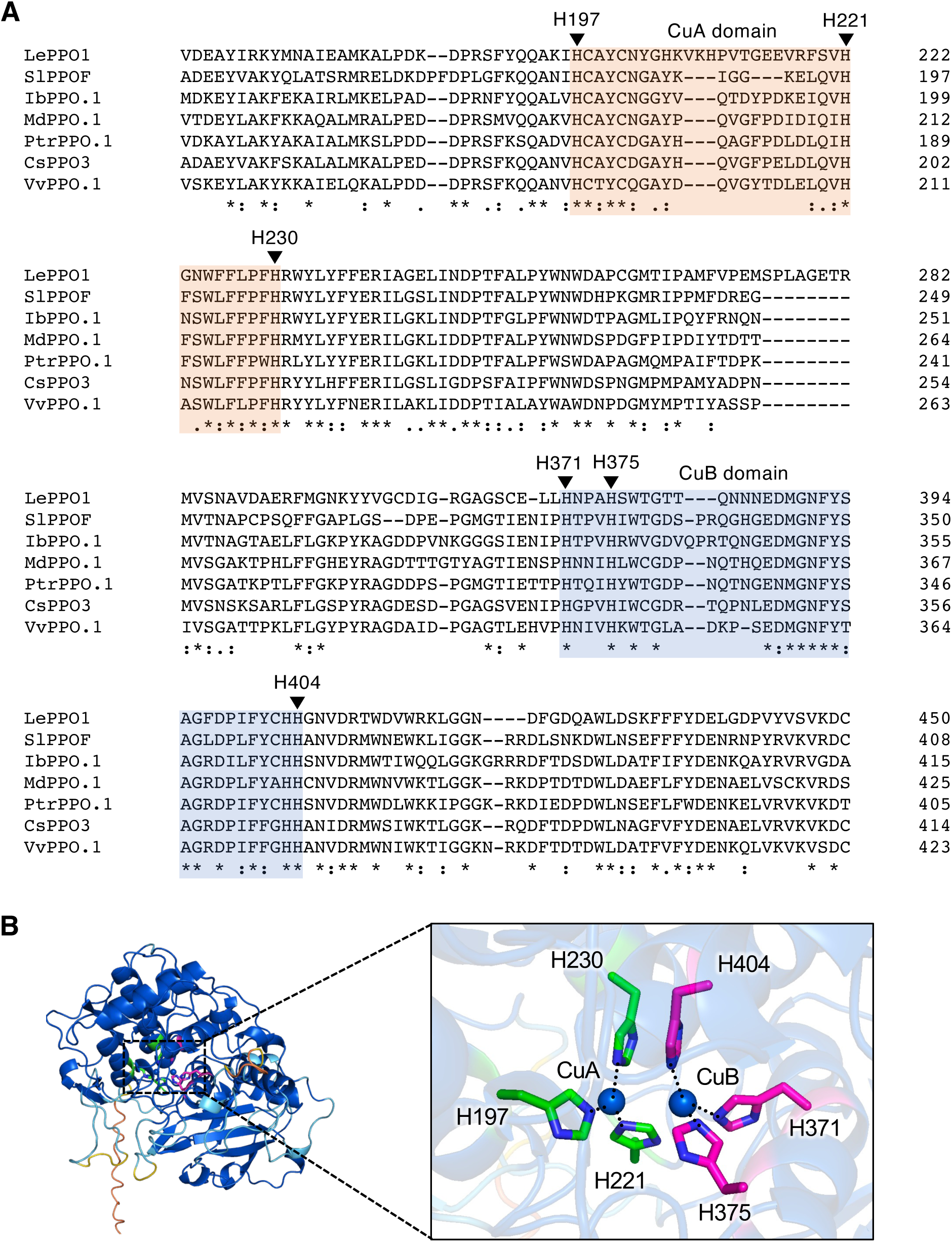
Conserved residues of plant PPOs and the predicted 3D-structure of LePPO1. A: Multiple sequence alignment of plant PPOs with LePPO1 and two copper ions (CuA and CuB) binding sites. Multiple sequence alignment was performed by Clustal Omega. Asterisks indicate consensus amino acid residues. CuA and CuB binding sites are highlighted by orange and blue boxes, respectively. His residues binding CuA and CuB are represented by triangle symbols. Cs, *Camellia sinensis*; Ib, *Ipomoea batatas*; Le, *Lithospermum erythrorhizon*; Md, *Malus domestica*; Ptr, *Populus trichocarpa*; Sl, *Solanum lycopersicum*; Vv, *Vitis vinifera*. B: The predicted structure of LePPO1 with two copper ions by AlphaFold 3 (pLDDT: orange, 0-50; yellow, 50-70; cyan, 70-90; blue, 90-100). His residues binding to CuA and CuB are shown as green color and magenta color, respectively. The structure was visualized by PyMOL.

Next, a neighbor-joining tree was constructed to evaluate the phylogenetic relationships among the five LePPOs and other reported plant PPOs. Some plant PPOs form family-specific clades, as seen in the Theaceae, Vitaceae, Salicaceae, Solanaceae, and Convolvulaceae families (Fig. 4). In Fabaceae and Rosaceae, however, PPO members formed two distinct clades. Interestingly, the LePPOs (Boraginaceae) were separated into three different clades. LePPO1, which is coordinately expressed with shikonin production, was classified in a clade near Solanaceae, along with LePPO2 and LePPO3 (Fig. 4). LePPO4 was classified in a clade close to Convolvulaceae, which is phylogenetically related to Boraginaceae and Solanaceae. LePPO5, on the other hand, was classified in a distant clade close to Vitaceae and Theaceae (Fig. 4). These results suggest that LePPO2, LePPO3, and LePPO4 have similar enzymatic activities to LePPO1 and are regulated by expression profiles, such as differential expression patterns or localizations. In contrast, LePPO5, which is 41.7% identical to LePPO1, has a different function than the other LePPOs.

**Figure 4.**
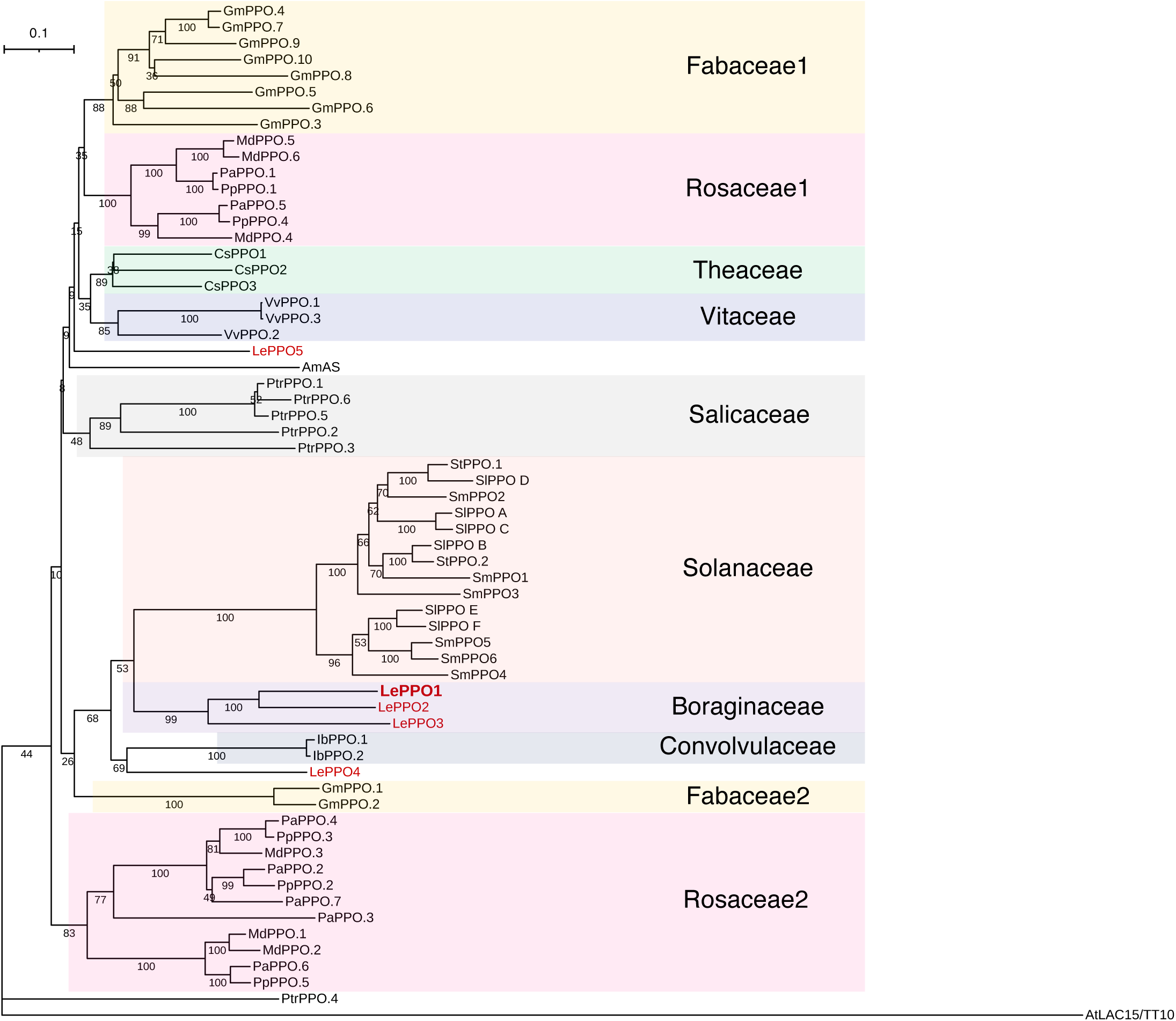
Phylogenetic relationship of LePPO1 and eudicot PPOs. The protein sequences were aligned by MUSCLE and a neighbor-joining tree was constructed using MEGA12 (v12.0.10) with the bootstrap method (1,000 replications). The five LePPOs are highlighted by red letters. AtLAC15/TT10 was used as an outgroup protein. The accession numbers of the protein sequences used in this study are listed in Supplementary Table S5. AS, aureusidin synthase; LAC, laccase; TT10, transparent testa10.

### Shikonin derivatives are strongly decreased in genome-edited hairy roots of *LePPO1*

To evaluate the role of LePPO1 in shikonin production in *L. erythrorhizon*, we generated genome-edited hairy roots of *LePPO1* (*LePPO1*-ge) via CRISPR/Cas9-mediated targeted mutagenesis. Four single-guide RNAs (sgRNAs) were designed in the CDS regions of *LePPO1* using the CRISPR-P 2.0 program (Fig. 5A). The genome editing constructs were then introduced into *L. erythrorhizon* via *Rhizobium rhizogenes* (strain ATCC15834). The stable transformants were then obtained as hairy roots. After genotyping, three independent *LePPO1*-ge lines (#112, #113, and #117) with different frameshift mutation patterns were selected (Supplemental Fig. S2 and Fig. 5B). AlphaFold 3 predicted that most mutation patterns would result in the loss of the CuB domain and three His residues. This occurred at a 100% ratio in all lines (Supplemental Fig. S2 and Fig. 5B). Line #113 of *LePPO1*-ge retained a partial CuB domain at a 70% frequency, however, the third His residue was lost, and the pLDDT score was below 50 in the AlphaFold 3 prediction (Fig. 5B). These results indicate that this mutant will not bind to CuB, nor will other genome-edited mutants.

**Figure 5.**
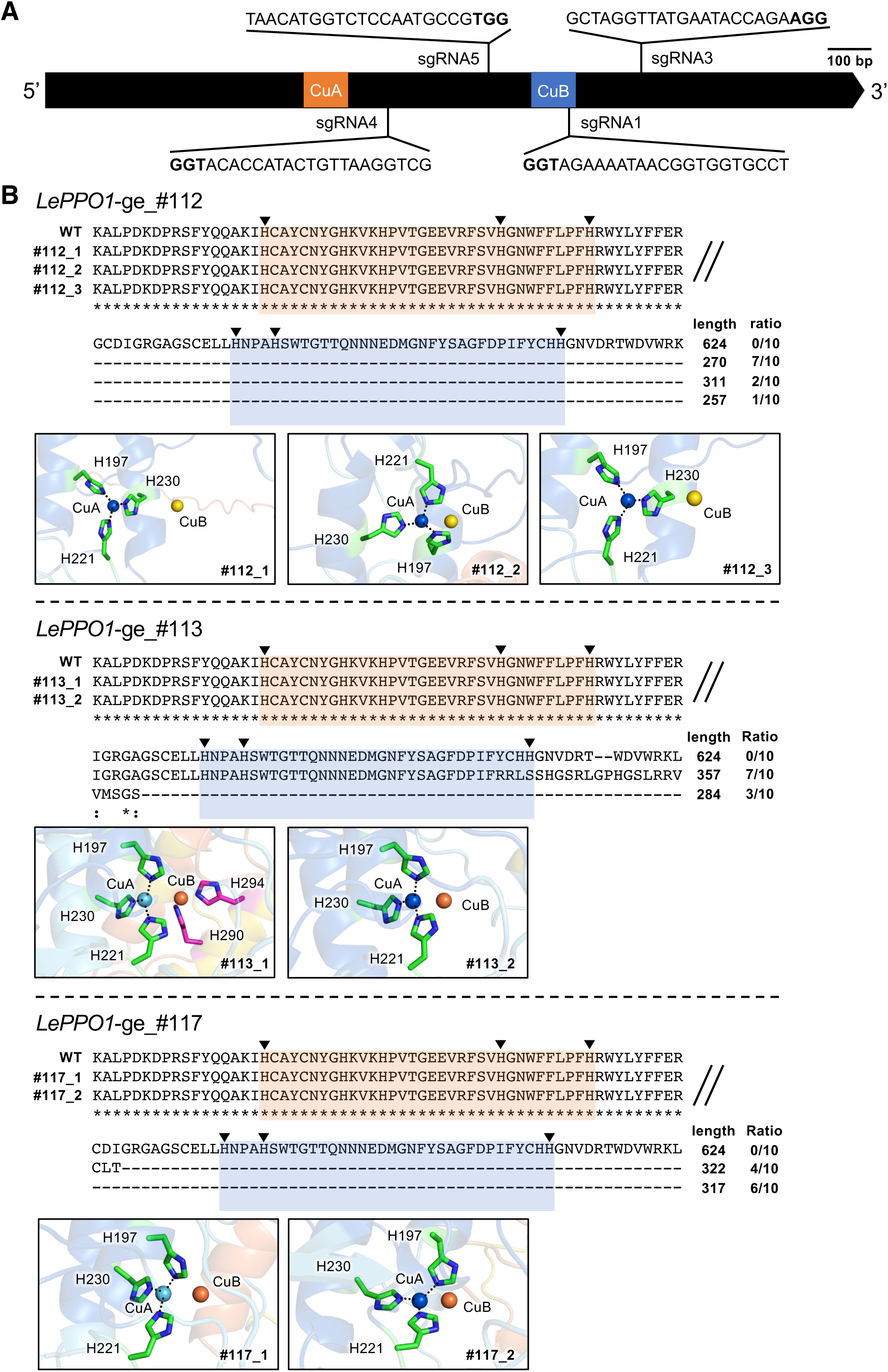
The designed sgRNAs and frameshift patterns of genome-edited lines. A: The location of four sgRNAs designed in *LePPO1*. B: Frameshift patterns, frameshift ratio, and the predicted structure of *LePPO1* genome-editing lines. Asterisks indicate consensus amino acids. CuA and CuB binding sites are highlighted by orange and blue colors, respectively. His residues required for binding CuA or CuB are represented by triangles. WT, wild type. Structures with two copper ions were predicted by AlphaFold 3 (pLDDT: orange, 0-50; yellow, 50-70; cyan, 70-90; blue, 90-100) and visualized by PyMOL. His residues binding CuA and CuB are shown as green color and magenta color, respectively.

Next, we extracted metabolites from *LePPO1*-ge lines cultured in M9 medium in the dark to induce shikonin derivative production. We analyzed the metabolites using the previously established HPLC method (Nakanishi et al. 2024) and compared them with those of a vector control line (Fig. 6A). *LePPO1*-ge lines produce several shikonin derivatives (naphthoquinone-type meroterpenes) and various specialized metabolites, including a shunt product (deoxyshikonofuran, a hydroquinone-type meroterpene) and two caffeic acid oligomers (lithospermic acid B, a tetramer, and rosmarinic acid, a dimer). Using the above HPLC conditions, all of these products can be detected with one injection. Figure 6C shows the contents of these metabolites, with shikonin derivatives shown as the sum of the detected derivatives. It should be noted that the total content of shikonin derivatives decreased significantly in all *LePPO1*-ge lines compared to the vector control (Supplemental Fig. S3 and Fig. 6C). In contrast, the contents of deoxyshikonofuran, lithospermic acid B, and rosmarinic acid did not change in these same lines (Supplemental Fig. S3 and Fig. 6C). These results demonstrate that *LePPO1* plays a crucial role in the shikonin-specific biosynthetic pathway and strongly suggest that the copper ion requirement for shikonin production is due to the reaction with PPO.

**Figure 6.**
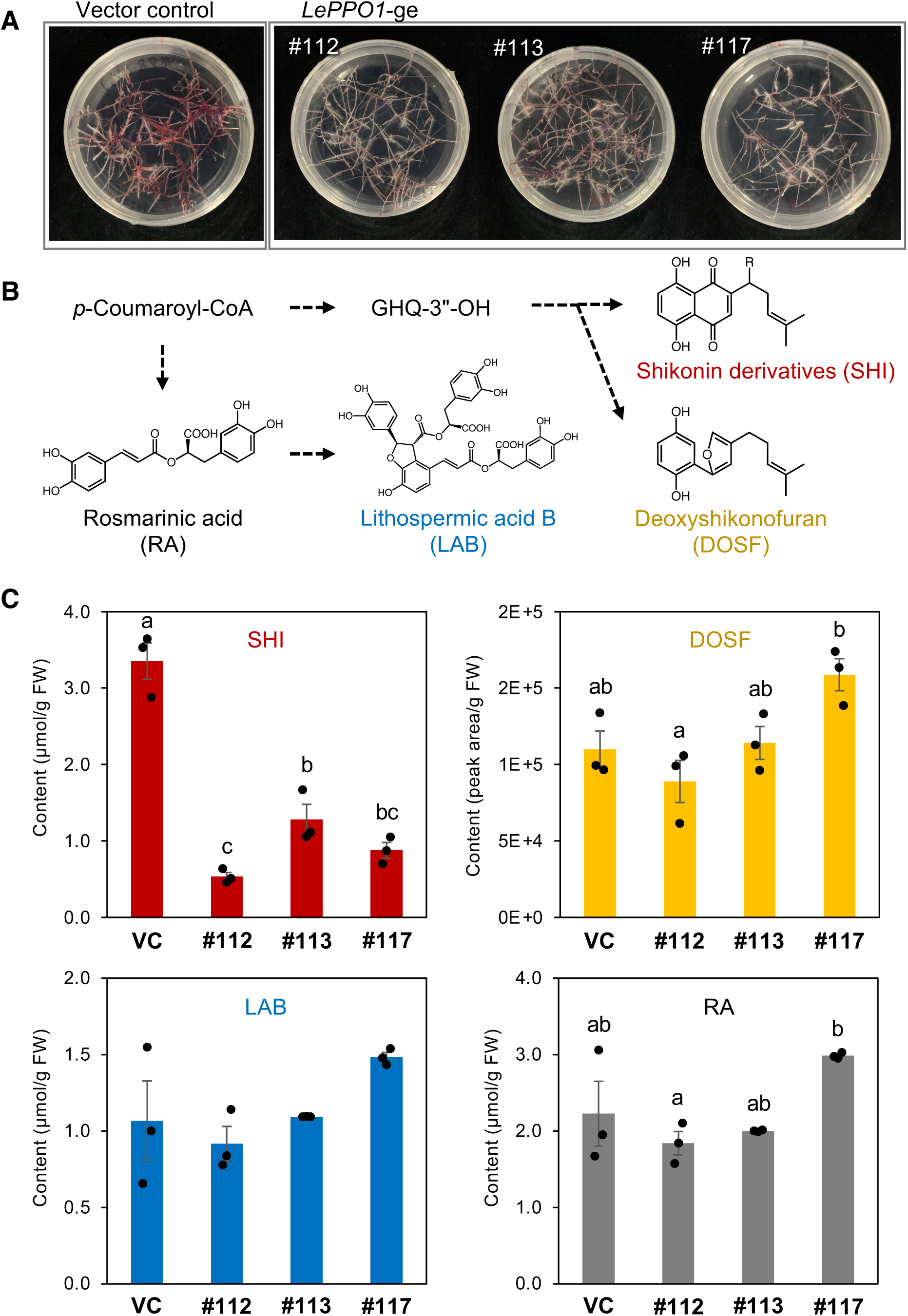
Metabolite analysis of *LePPO1* genome-edited lines. A: The LePPO1 genome-edited hairy roots cultured in M9 medium for 26 days in the dark. B: The overview of four endproducts produced by hairy roots of *L. erythrorhizon*. C: Content of shikonin derivatives (SHI), deoxyshikonofuran (DOSF), lithospermic acid B (LAB), and rosmarinic acid (RA) in three biologically independent lines of *LePPO1*-ge and vector control. Bars are represented as mean ± SE of technically independent cultures (*n* = 3). Different letters indicate significant difference by Tukey’s honestly significant difference test *(P* < 0.05).

## Discussion

Shikonin derivatives are red 1,4-naphthoquinone compounds that exhibit diverse pharmacological properties. The roots of *L. erythrorhizon*, which contain shikonin derivatives, have been used as a crude drug and as an ingredient in pharmaceutical products. They have also been used as natural dyes to stain traditional garments, such as kimonos, throughout Japan’s long history. Due to their benefits and safety, *L. erythrorhizon* roots and shikonin derivatives are also used in cosmetics. Due to their commercial value and scientific interest, the biosynthetic pathway of shikonin has been actively studied for the past half century. In recent years, many biosynthetic genes have been identified, however, the formation of the naphthalene ring remains unclear. It is only expected that some oxidoreductases are involved, however, it is still under discussion whether naphthalene formation is a single reaction or a series of successive enzyme reactions. Additionally, the reason why the copper ion is essential for shikonin production remains elusive. In this study, we identified *LePPO1*, one of five copper-dependent *polyphenol oxidase* genes in *L. erythrorhizon* that shows clear expression correlated with shikonin production. We have demonstrated via genome editing techniques that *LePPO1* plays a crucial role in shikonin production.

Various differential screening methods have been applied to the *L. erythrorhizon* cell culture system to identify target genes involved in shikonin biosynthesis (Yazaki et al. 2001). Among these methods, the PCR-select subtraction method appears to be the most effective for identifying target genes, as has been reported in various organisms, including plant species (Kajimoto et al. 2003, Yamazaki et al. 2008, Sahu and Shaw 2009, Bunsupa et al. 2011). In our experiment, the PCR-select subtraction method narrowed down many biosynthetic genes that had already been identified, as well as many fragments of uncharacterized *PPO* genes in *L. erythrorhizon* cells under conditions that induce shikonin production (Fig. 1). Transcriptome analysis identified five *LePPO* genes expressed in cultured *L. erythrorhizon* cells. Narrowing down the robust candidates revealed that *LePPO1* showed an obvious strong correlation (Fig. 2). *LePPO5*, the most divergent *PPO* among the five *LePPOs*, was the most highly expressed and showed ubiquitous expression throughout the culture period and in plant organs. This suggests the possibility of its involvement in common metabolism. Additionally, many clones predicted to be involved in transporters and other metabolic processes were highlighted. Future studies will use the data from PCR subtraction to elucidate the excretion mechanism of shikonin derivatives (Tatsumi et al. 2016, Kiyoto et al. 2022, Tatsumi et al. 2023, Ichino et al. 2025).

Multiple alignment analysis of PPO protein sequences, including LePPO1 and previously reported plant PPOs, revealed that the conserved sequences CuA and CuB are present in LePPO1 (Fig. 3). However, several unique amino acid residues were found in the CuA domain of LePPO1. For example, SlPPOF, which is found in tomato and has unique amino acid residues in the CuA domain, catalyzes the conversion of eriodictyol to luteolin in flavone biosynthesis (Wei et al. 2022). The unique amino acid residues of LePPO1 may be related to its substrate recognition and/or specificity in specialized metabolism.

The diversity of enzymatic reactions is a key feature of PPO. The browning effect in crops is caused by PPO, which oxidizes phenolic substances such as chlorogenic acid (Tran et al. 2012). The PPO in snapdragons and *Marchantia polymorpha* catalyze the formation of the furan ring of the chalcone (Nakayama et al. 2000, Nakayama et al. 2001, Furudate et al. 2023). Additionally, oxidative polymerization of flavonoids leading to proanthocyanidin formation is catalyzed by a PPO (Pourcel et al. 2005). CsPPO3 in *Camellia sinensis* is highly expressed in the dark and catalyzes the conversion of catechins to theaflavins (Yu et al. 2020). VvPPO2 in Shine Muscat has been characterized as a member involved in skin browning (Katayama-Ikegami et al. 2017). These reports suggest high flexibility in PPO enzymatic reactions (Sullivan 2015). Since LePPO5 is classified in a clade distant from the other LePPOs and close to CsPPO3 and VvPPO2, it may be responsible for a common metabolism unlike shikonin biosynthesis (Fig. 4).

Previously, we proposed that single-exon genes might be involved in secondary metabolism (Kusano et al. 2019). Within *L. erythrorhizon*, the enzyme responsible for the coupling of the aromatic substrate (PHB) and the isoprenoid precursor (GPP), *LePGT*, is intronless. However, other *L. erythrorhizon prenyltransferases* involved in ubiquinone biosynthesis include introns. Similarly, the *4-coumaroyl-CoA ligase* genes responsible for shikonin formation, *Le4CL3* and *Le4CL4*, are intronless. In contrast, the *4CLs* involved in lignin formation have introns (Nakanishi et al. 2024). The *LePPO1* data from this study support the hypothesis that single-exon genes tend to be involved in secondary metabolism (Fig. 5A).

We previously demonstrated that the genome editing of *Le4CL3* or *Le4CL4* in hairy roots suppressed the production of shikonin and echinofuran (benzoquinone) derivatives (Nakanishi et al. 2024). This is because 4CL is responsible for an upstream biosynthetic reaction of both metabolites. In contrast, genome editing of *LePPO1* did not affect the content of the echinofuran derivative and the caffeic acid derivatives, such as lithospermic acid B. However, shikonin production was clearly suppressed (Fig. 5 and 6). These results indicate that LePPO1 is responsible for downstream reaction steps specific for shikonin biosynthesis, i.e., those after the hydroquinone/benzoquinone metabolism branch point (Fig. 6B). Thus, we hypothesized that LePPO1 is involved in the naphthalene ring formation step. Using HPLC analysis, we attempted to detect the generation of a new metabolite in genome-edited mutants of *LePPO1* as a putative biosynthetic intermediate, but we could not detect any new peaks compared to the control hairy roots. Identifying the direct substrate of LePPO1 and its reaction mechanism will be the focus of further studies.

In summary, we demonstrated through omics data analysis and a reverse genetic approach that dicopper-dependent LePPO1 participates in shikonin production. Shikonin derivatives are being investigated for their potential benefits to humans, including their pharmacological properties (Han et al. 2025). The findings of this study will further our understanding of the biosynthesis of shikonin and other 1,4-naphthoquinone derivatives in plants.

## Materials and Methods

### Chemicals and reagents

*p*-Hydroxybenzoic acid, *p*-coumaric acid, rosmarinic acid, and lithospermic acid B (salvianolic acid B) were purchased from Tokyo Chemical Industry (Tokyo, Japan), FUJIFILM Wako (Tokyo, Japan), and MedChemExpress (Monmouth Junction, NJ, USA), respectively. *β*-Hydroxyisovarelylshikonin, acetylshikonin, isobutyrylshikonin, and isovarelylshikonin were purchased from Nagara Science (Gifu, Japan). GHQ-3′′-OH was synthesized according to our previous report (Oshikiri et al., 2020). All antibiotics were purchased from FUJIFILM Wako.

### Plant materials and growth conditions

Cultured *L. erythrorhizon* (strain TOM) cells have been subcultured in Linsmaier and Skoog (LS) medium containing 3% (w/v) sucrose, 10^-6^ M indole-3-acetic acid (IAA), and 10^-5^ M kinetin at pH 6.5 in darkness at 25 °C (summarized in Yazaki 2017). Subcultures were repeated approximately every two months until they were used for PCR-select subtraction. Sterile shoot cultures of *L. erythrorhizon* were grown on a 1/2 Murashige and Skoog (MS) agar medium containing 1% sucrose by weight at pH 5.6 under 16-hour light and 8-hour dark conditions. Subcultures were repeated every three months until they were used for hairy root induction. *L. erythrorhizon* hairy roots were cultured in 1/2 MS agar medium containing 1% sucrose and 20 mg/L meropenem at pH 5.6 in darkness at 23 °C. Subcultures were repeated once every three months until they were used for genotyping and metabolite analysis.

### Total RNA extraction and cDNA synthesis

Total RNA was extracted using the RNeasy Plant Mini Kit (Qiagen, Venlo, Netherlands) according to the manufacturer’s instructions. cDNA was synthesized using the DNA-free™ Kit (Invitrogen, Waltham, MA) and SuperScript™ III Reverse Transcriptase (Invitrogen). The RNA and cDNA samples were stored at -80 °C.

### PCR-select cDNA subtraction and cDNA cloning

Total RNA was prepared from TOM cells cultured in an M9 medium containing 3% sucrose, 10^-6^ M IAA, and 10^-5^ M kinetin at pH 6.5. The cells were cultured for 12 days at 25 °C under light or in the dark at 80 rpm. PCR-select subtraction was performed between the cDNAs from the light and dark conditions (Diatchenko et al. 1996, Gurskaya et al. 1996). According to the PCR-select subtraction library, *LePPO1*-specific primers were designed (Supplementary Table S3). cDNA cloning of *LePPO1* was performed by the 5’/3’ RACE method using a GeneRacer Kit (Invitrogen) according to the instruction manual with gene-specific primers and the cDNA from the dark sample as a template. Full-length *LePPO1* was amplified, subcloned into a pT7 Blue T-vector via TA cloning, and sequenced.

### Transcriptome analysis

RNA-seq data for *L. erythrorhizon* were obtained from the National Center for Biotechnology Information Sequence Read Archive (SRA; https://www.ncbi.nlm.nih.gov/sra). Their SRA codes are listed in Supplementary Table S1. The RNA-seq data were trimmed using Trimmomatic (version 0.39) with the ILLUMINACLIP option (Bolger et al. 2014). The trimmed reads were then mapped to the genome assembly GCA_912073615.1 (Tang 2021) using HiSAT2 (v 2.2.1_1) (Kim et al. 2019). StringTie (v 3.0.0) was used to calculate read counts and transcripts per million (TPM) values (Pertea et al. 2015). The TPM values were transformed to log2 (TPM+1), normalized by Z-score, and a heatmap was generated using the *pheatmap* package in RStudio (v 2024.12.1+563).

### Gene expression analysis

Total RNA was prepared from each plant tissue (flower, bud, upper leaf, lower leaf, stem, main root, lateral root, root bark, and peeled root) of *L. erythrorhizon* and used for cDNA synthesis. The cDNAs were then used for real-time quantitative PCR (RT-qPCR) analysis. RT-qPCR was performed using Platinum SYBR Green qPCR SuperMix-UDG (Invitrogen) and gene-specific primers (Supplementary Table S3) with a C1000 Touch Thermal Cycler and CFX96 Deep Well Real-Time System (Bio-Rad, Hercules, CA, USA). The PCR reaction was carried out at 95 °C for 1 min, followed by 40 cycles of 95 °C for 10 sec, 57 °C for 15 sec, 72 °C for 30 sec, and 82 °C for 5 sec. A melt curve was generated by heating the samples from 65 °C to 95 °C at 0.5 °C increments, each lasting 5 seconds, while acquiring data in real time. The *L. erythrorhizon actin* gene (*LeACT*) was used as the reference gene (Supplementary Table S3).

### Multiple sequence alignment and phylogenetic analysis

We searched the protein sequences of plant PPOs and laccases (LACs) in the UniProt database (https://www.uniprot.org/) and the NCBI database. Plant PPOs ranging in length from 450 to 650 amino acids and plant LACs were retrieved from NCBI. Multiple sequence alignment of LePPOs and plant PPOs was performed using Clustal Omega (https://www.ebi.ac.uk/Tools/msa/clustalo/) with default settings (Larkin et al. 2007). The GenBank accession numbers of the plant PPOs and LACs used in this study are listed in Supplementary Tables S4 and S5. The protein sequences were aligned using MUSCLE with the default options (Edgar 2004). A neighbor-joining tree was constructed using the bootstrap method (1,000 replicates) with MEGA12 (v 12.0.10) (Kumar et al. 2024). AtLAC15/TT10 was used as the outgroup for the plant PPOs. The phylogenetic tree was visualized using iTOL (https://itol.embl.de/).

### Generation and culturing of genome-edited hairy roots

The generation of the genome-edited hairy roots of *L. erythrorhizon* was performed using the previously described method (Nakanishi et al. 2024). Four sgRNAs targeting the CDS region of *LePPO1* were designed using the CRISPR-P 2.0 program, as previously described (Liu et al. 2017). The blastn-short program was used to identify off-targets for all selected sgRNAs. PCR primers including these sgRNAs were designed, and the sgRNA fragments were amplified. The sgRNA fragments ligated using the Golden Gate cloning method (Engler et al. 2008) were inserted into the BsaI site of pMgP237-2A-GFP (Ueta et al. 2017, Nakayasu et al. 2018). After Bsa I treatment, the construct was transformed into the *Escherichia coli* DH5α strain. Purified plasmid DNA was then transformed into a competent *Rhizobium rhizogenes* strain (ATCC15834) by electroporation using a MicroPulser electroporator (Bio-Rad, Hercules, CA, USA). Transformation into leaf disks of *L. erythrorhizon* was then performed (Tatsumi et al. 2020). Genomic DNA (gDNA) was extracted from hairy roots and used for genotyping to confirm mutagenesis. All of the original primers used in this study are summarized in Supplementary Table S3. To determine the mutation patterns of the genome-edited lines, the amplified fragments that annealed with an A residue using a 10x A-attachment mix (TOYOBO) were inserted into a pGEM-T Easy vector (Promega, Madison, WI, USA) according to the manufacturer’s instructions. Each vector was then transformed into *E. coli* DH5α competent cells, and ten positive colonies were obtained from each sample by blue/white screening with ampicillin as the selection marker. All plasmid DNAs were extracted using a modified protocol of the FastGene Plasmid Mini Kit (NIPPON Genetics, Tokyo, Japan) and sequenced using T7 and SP6 primers (Shimizu et al. 2023). Amino acid sequences after frameshift were predicted using the translation option in Benchling (https://www.benchling.com/). The 3D structure of LePPO1 and its mutants with two copper ions was predicted using AlphaFold 3 (pLDDT: orange, 0–50; yellow, 50–70; cyan, 70–90; blue, 90– 100) (Abramson et al. 2024) and visualized using PyMOL. The generated hairy roots were subcultured in 1/2 MS agar medium containing meropenem.

### Metabolite analysis of hairy roots

Hairy roots were cultured in an M9 agar medium in the dark at 23 °C for 26 days and were used to analyze the knock-out ratio and metabolites. Metabolite extraction from the hairy root samples and high-performance liquid chromatography (HPLC) analysis were performed using our previous method (Nakanishi et al. 2024). Shortly after extraction, the methanol extracts of genome-edited hairy roots and vector controls were analyzed by HPLC. The content of shikonin derivatives, echinofuran derivatives, lithospermic acid B (LAB), and rosmarinic acid were calculated using the standard curve method from the peak area at 280 nm for the echinofuran derivative, 290 nm for LAB, 330 nm for rosmarinic acid, and 520 nm for the shikonin derivative.

### Data analysis

The data are presented as bars representing the mean and points representing single values, with error bars representing the standard deviation (SD) or standard error (SE) of technical replicates or biologically independent lines (n ≥ 3). One-way analysis of variance (ANOVA) was performed, followed by the Tukey’s honestly significant difference (HSD) test for multiple comparisons.

## Accession numbers

The accession numbers of the RNA-seq data, genes, and proteins used in this study are described in Supplementary Tables S1, S2, S4 and S5 and are available in the Sequence Read Archive (SRA) or GenBank data libraries.

## Data availability

All generated and analyzed data from this paper are included in the published article and Supplemental Data.

## Acknowledgments

We thank Mr. Yuji Sakai of Kazusa DNA Research Institute for support of sequencing; Dr. Hirobumi Yamamoto for providing the benzoquinone derivatives; Ms. Saw Yu Yu Hnin of Toyama University for technical support of 3D-structure prediction; Dr. Kanako Sasaki and Ms. Eiko Moriyoshi of Kyoto University for technical support of genome editing experiments; Dr. Noboru Onishi of Kirin Holdings Company, Ltd. for providing sterile shoot cultures of *L. erythrorhizon*; Amato Pharmaceutical Products, Ltd. for providing seeds and plant tissue samples of *L. erythrorhizon*. Part of this study was performed using the DASH/FBAS facilities at RISH, Kyoto University.

## Author contributions

K.N., T.I., Ka.T., H.L., A.S., K.T., and Ka.Y. conceived and designed the study; Ka.O., R.M., H.S., N.S., D.S., Ke.O., A.S., Ko.T., and Ka.Y. supervised the experiments; K.N., Y.T., Ky.Y, M.Y., K.M., T.O., and K.S. performed the experiments; B.W. synthesized GHQ-3’’-OH; K.N. and Ka.Y. wrote the manuscript; all authors discussed the results, reviewed the article, and approved the manuscript.

## Supporting Information

**Supplemental Figure S1.** Phylogenetic relationship of plant polyphenol oxidases (PPOs) and laccases (LACs). The protein sequences were aligned by MUSCLE and Neighbor-joining tree was constructed by MEGA12 (v12.0.10) with bootstrap method (1,000 replications). LePPO1 is represented by red letters. The accession numbers of protein sequences used in this analysis are described in supplementary Table S4. Am, *Antirrhinum majus*; Cs, *Camellia sinensis*; Gm, *Glycine max*; Ib, *Ipomoea batatas*; Le, *Lithospermum erythrorhizon*; Md, *Malus domestica*; Os, *Oryza sativa*; Pa, *Prunus armeniaca*; Pp, *Prunus persica*; Ptr, *Populus trichocarpa*; Sl, *Solanum lycopersicum*; Sm, *Solanum melongena*; St, *Solanum tuberosum*; Vv, *Vitis vinifera*; Zm, *Zea mays*. AS, aureusidin synthase; TT10, transparent testa10.

**Supplemental Figure S2.** Mutation patterns of *LePPO1* genome-edited lines. The mutations on sgRNAs are shown as red color. WT, wild type.

**Supplemental Figure S3.** HPLC chromatograms of *LePPO1* genome-edited lines and vector control (λ=280 nm). PHB-OG, *p*-hydroxybenzoic acid-*O*-glucoside; PCA, *p*-coumaric acid; RA, rosmarinic acid; LAB, lithospermic acid B; GHQ, geranylhydroquinone; DHSF, dihydroshikonofuran; GBA, *m*-geranyl-*p*-hydroxybenzoic acid; HIVS, *β*-hydroxyisovarelylshikonin; AS, acetylshikonin; EFB, echinofuran B; DOSF, deoxyshikonofuran; IBS, isobutyrylshikonin; IVS, isovarelylshikonin.

**Supplementary Table S1.** RNA-seq data used in transcriptome analysis

**Supplementary Table S2.** Genes used in expression profile analysis

**Supplementary Table S3.** Primers used in this study

**Supplementary Table S4.** Plant PPOs and LACs used in Supplementary Figure S1

**Supplementary Table S5.** Plant PPOs used in this study

## References

Abramson, J., Adler, J., Dunger, J., Evans, R., Green, T., Pritzel, A., et al. (2024) Accurate structure prediction of biomolecular interactions with AlphaFold 3. Nature 630: 493–500

Alonso, A.V., Naranjo, H.D., Rat, A., Rodić, N., Nannou, C.I., Lambropoulou, D.A., et al. (2022) Root-associated bacteria modulate the specialised metabolome of *Lithospermum officinale* L.. Front. Plant Sci. 13: 908669

Auber, R.P., Suttiyut, T., McCoy, R.M., Ghaste, M., Crook, J.W., Pendleton, A.L., et al. (2020) Hybrid de novo genome assembly of red gromwell (*Lithospermum erythrorhizon*) reveals evolutionary insight into shikonin biosynthesis. Hortic. Res. 7: 82

Bai, Y., Ali, S., Liu, S., Zhou, J., Tang, Y. (2023) Characterization of plant laccase genes and their functions. Gene 852: 147060

Bolger, A.M., Lohse, M., Usadel, B. (2014) Genome analysis Trimmomatic : a flexible trimmer for Illumina sequence data. Bioinformatics 30: 2114–2120

Brigham, L.A., Michaels, P.J., Flores, H.E. (1999) Cell-specific production and antimicrobial activity of naphthoquinones in roots of *Lithospermum erythrorhizon*. Plant Physiol. 119: 417–428

Bunsupa, S., Okada, T., Saito, K., Yamazaki, M. (2011) An acyltransferase-like gene obtained by differential gene expression profiles of quinolizidine alkaloid-producing and nonproducing cultivars of *Lupinus angustifolius*. Plant Biotechnol. 28: 89–94

Chen, Q., Ding, Y., Li, Z., Chen, X., Fazal, A., Zhang, Y., et al. (2025) Comprehensive analysis of the antibacterial activity of 5,8-dihydroxy-1.4-naphthoquinone derivatives against methicillin-resistant *Staphylococcus aureus*. Chin. J. Nat. Med. 23: 604–613

Diatchenko, L., Laut, Y.C., Campbellt, A.P., Chenchik, A., Moqadam, F., Huang, B., et al. (1996) Suppression subtractive hybridization: A method for generating differentially regulated or tissue-specific cDNA probes and libraries. Proc. Natl. Acad. Sci. U.S.A. 93: 6025–6030

Edgar, R.C. (2004) MUSCLE: Multiple sequence alignment with high accuracy and high throughput. Nucleic Acids Res. 32: 1792–1797

Engler, C., Kandzia, R., Marillonnet, S. (2008) A one pot, one step, precision cloning method with high throughput capability. PLoS ONE 3: e3647

Fu, J., Lu, G., Yang, M., Zhao, H., Jie, W., Fazal, A., et al. (2021) Cloning and functional analysis of *EpGHQH1* in shikonin production of *Echium plantagineum*. Plant Cell, Tissue Organ Cult. doi: 10.1007/s11240-020-01976-2

Fukui, H., Yazaki, K., Tabata, M. (1984) Two phenolic acids from *Lithospermum erythrorhizon* cell suspension cultures. Phytochemistry 23: 2398–2399

Fujita, Y., Hara, Y., Suga, C., Morimoto, T. (1981) Production of Shikonin Derivatives by Cell Suspension Cultures of *Lithospermum erythrorhizon* - II. A new Medium for the Production of Shikonin Derivatives. Plant Cell Rep. 1: 61–63

Furudate, H., Manabe, M., Oshikiri, H., Matsushita, A., Watanabe, B., Waki, T., et al. (2023) A polyphenol oxidase catalyzes aurone synthesis in *Marchantia polymorpha*. Plant Cell Physiol. 64: 637–645

Guo, C., He, J., Song, X., Tan, L., Wang, M., Jiang, P., et al. (2019) Pharmacological properties and derivatives of shikonin — A review in recent years. Pharmacol. Res. 149: 104463

Gurskaya, N.G., Diatchenko, L., Chenchik, A., Siebert, P.D., Khaspekov, G.L., Lukyanov, K.A., et al. (1996) Equalizing cDNA Subtraction Based on Selective Suppression of Polymerase Chain Reaction : Cloning of Jurkat Cell Transcripts Induced by Phytohemaglutinin and Phorbol 12-Myristate 13-Acetate. Anal. Biochem. 240: 90–97

Han, H., Wen, Z., Yang, M., Wang, C., Ma, Y., Chen, Q., et al. (2025) Shikonin Derivative Suppresses Colorectal Cancer Cells Growth via Reactive Oxygen Species-Mediated Mitochondrial Apoptosis and PI3K / AKT Pathway. Chem. Biodivers. 22: 1–12

Ichino, T., Tatsumi, K., Munakata, Y., Tsuboyama, A., Moriyoshi, E., Nakayasu, M. et al. (2025) Characterization of two G-type half-size ABC transporter genes from the lipid-secreting medicinal plant *Lithospermum erythrorhizon*. J. Plant Res. 10.1007/s10265-025-01651-7

Ito, E., Munakata, R., Yazaki, K. (2023) Letter to the Editor: Gromwell, a Purple Link between Traditional Japanese Culture and Plant Science. Plant Cell Physiol. 64: 567–570

Izuishi, Y., Isaka, N., Li, H., Nakanishi, K., Kageyama, J., Ishikawa, K., et al. (2020) Apple latent spherical virus (ALSV)-induced gene silencing in a medicinal plant, *Lithospermum erythrorhizon*. Sci. Rep. 10: 1–9

Jin, Z., Du, X., Xu, Y., Deng, Y., Liu, M., Zhao, Y., et al. (2020a) Structure of Mpro from SARS-CoV-2 and discovery of its inhibitors. Nature 582: 289–293

Jin, Z., Zhao, Y., Sun, Y., Zhang, B., Wang, H., Wu, Y., et al. (2020b) Structural basis for the inhibition of SARS-CoV-2 main protease by antineoplastic drug carmofur. Nat. Struct. Mol. Biol. 27: 529–532

Kajimoto, K., Daikoku, T., Kita, F., Yamazaki, N., Kataoka, M., Baba, Y., et al. (2003) PCR-select subtraction for characterization of messages differentially expressed in brown compared with white adipose tissue. Mol. Genet. Metab. 80: 255–261

Katayama-ikegami, A., Suehiro, Y., Katayama, T., Jindo, K., Esumi, T. (2017) Recombinant expression, puri fi cation, and characterization of polyphenol oxidase 2 (*VvPPO2*) from “Shine Muscat” (*Vitis labruscana* Bailey × *Vitis vinifera* L.). Biosci. Biotechnol. Biochem. 81: 2330–2338

Kim, D., Paggi, J.M., Park, C., Bennett, C., Salzberg, S.L. (2019) Graph-based genome alignment and genotyping with HISAT2 and HISAT-genotype. Nat. Biotechnol. 37: 907–915

Kiyoto, S., Ichino, T., Awano, T., Yazaki, K. (2022) Improved chemical fixation of lipid-secreting plant cells for transmission electron microscopy. Microscopy 71: 206–213

Kumar, S., Stecher, G., Suleski, M., Sanderford, M., Sharma, S., Tamura, K. (2024) MEGA12 : Molecular Evolutionary Genetic Analysis Version 12 for Adaptive and Green Computing. Mol. Biol. Evol. 41: 1–9

Kusano, H., Li, H., Minami, H., Kato, Y., Tabata, H., Yazaki, K. (2019) Evolutionary developments in plant specialized metabolism, exemplified by two transferase families. Front. Plant Sci. 10: 794

Larkin, M.A., Blackshields, G., Brown, N.P., Chenna, R., Mcgettigan, P.A., Mcwilliam, H., et al. (2007) Clustal W and Clustal X version 2.0. Bioinformatics 23: 2947–2948

Li, H., Matsuda, H., Tsuboyama, A., Munakata, R., Sugiyama, A., Yazaki, K. (2022) Inventory of ATP-binding cassette proteins in *Lithospermum erythrorhizon* as a model plant producing divergent secondary metabolites. DNA Res. 29: 1–12

Liang, M., Davis, E., Gardner, D., Cai, X., Wu, Y. (2006) Involvement of AtLAC15 in lignin synthesis in seeds and in root elongation of Arabidopsis. Planta 224: 1185–1196

Liao, Z., Chen, R., Chen, M., Yang, Y., Fu, Y., Zhang, Q., et al. (2006) Molecular Cloning and Characterization of the Polyphenol Oxidase Gene from Sweetpotato 1. Mol. Biol. 40: 907– 913

Liu, H., Ding, Y., Zhou, Y., Jin, W., Xie, K., Chen, L.L. (2017) CRISPR-P 2.0: An Improved CRISPR-Cas9 Tool for Genome Editing in Plants. Mol. Plant. 10: 530–532

Mayer, A.M. (2006) Polyphenol oxidases in plants and fungi : Going placesA review. Phytochemistry 67: 2318–2331

Nakanishi, K., Li, H., Ichino, T., Tatsumi, K., Osakabe, K., Watanabe, B., et al. (2024) Peroxisomal 4-coumaroyl-CoA ligases participate in shikonin production in *Lithospermum erythrorhizon*. Plant Physiol. 195: 2843–2859

Nakayama, T., Sato, T., Kikuchi, S., Fukui, Y., Ueda, T., Nakao, M., et al. (2000) Aureusidin Synthase : A Polyphenol Oxidase Homolog Responsible for Flower Coloration. Science 290: 1163–1166

Nakayama, T., Sato, T., Fukui, Y., Yonekura-Sakakibara, K., Hayashi, H., Tanaka, Y., et al. (2001) Specificity analysis and mechanism of aurone synthesis catalyzed by aureusidin synthase, a polyphenol oxidase homolog responsible for flower coloration. FEBS Lett. 499: 107–111

Nakayasu, M., Akiyama, R., Lee, H.J., Osakabe, K., Osakabe, Y., Watanabe, B., et al. (2018) Generation of α-solanine-free hairy roots of potato by CRISPR/Cas9 mediated genome editing of the *St16DOX* gene. Plant Physiol. Biochem. 131: 70–77

Okada, T. and Watanabe, K. (2024) The complete chloroplast genome sequence of *Lithospermum erythrorhizon*: Insights into the phylogenetic relationship among Boraginaceae species and the material linages of purple gromwells. Plant Gene 37: 100447

Oshikiri, H., Li, H., Manabe, M., Yamamoto, H., Yazaki, K., Takanashi, K. (2024) Comparative Analysis of Shikonin and Alkannin Acyltransferases Reveals Their Functional Conservation in Boraginaceae. Plant Cell Physiol. 65: 362–371

Oshikiri, H., Watanabe, B., Yamamoto, H., Yazaki, K., Takanashi, K. (2020) Two bahd acyltransferases catalyze the last step in the shikonin/alkannin biosynthetic pathway. Plant Physiol. 184: 753–761

Ozaki, Y., Sakaguchi, I., Tujimura, M., Ikeda, N., Nakayama, M., Kato, Y., Suzuki, H., Satake, M. (1998) Study of the Accelerating Effect of Shikonin and Alkanine on the Proliferation of Granulation Tissue in Rat. Biol. Pharm. Bull. 21: 366–370

Pertea, M., Pertea, G.M., Antonescu, C.M., Chang, T., Mendell, J.T., Salzberg, S.L. (2015) StringTie enables improved reconstruction of a transcriptome from RNA-seq reads. Nat. Biotechnol. 33: 290–295

Pourcel, L., Routaboul, J.-M., Kerhoas, L., Caboche, M., Lepiniec, L., Debeaujon, I. (2005) *TRANSPARENT TESTA10* Encodes a Laccase-Like Enzyme Involved in Oxidative Polymerization of Flavonoids in *Arabidopsis* Seed Coat. Plant Cell 17: 2966–2980

Rat, A., Koletti, A.E., Papageorgiou, V.P., Willems, A., Assimopoulou, A.N. (2023) Bacterial responses to plant antimicrobials : the case of alkannin and shikonin derivatives. Front. Pharmacol. 14: 1244270

Sahu, B.B., Shaw, B.P. (2009) Isolation, identification and expression analysis of salt-induced genes in *Suaeda maritima*, a natural halophyte, using PCR-based suppression subtractive hybridization. BMC Plant Biol. 9: 1–25

Sajjad, N., Ahmad, M.S., Mahmood, R.T., Tariq, M., Asad, M.J., Irum, S., et al. (2023) Purification and characterization of novel isoforms of the polyphenol oxidase from *Malus domestica* fruit pulp. PLoS ONE 18: e0276041

Sankawa, U., Ebizuka, Y., Miyazaki, T., Isomura, Y., Hideaki, O., Shibata, S., et al. (1977) Antitumor Activity of Shikonin and Its Derivatives. Chem. Pharm. Bull. 25: 2392–2395

Shimizu, K., Akiyama, R., Okamura, Y., Ogawa, C., Masuda, Y., Sakata, I., et al (2023) Solanoeclepin B, a hatching factor for potato cyst nematode. Sci. Adv. 9: 1–11

Song, W., Zhuang, Y., Liu, T. (2020) Potential role of two cytochrome P450s obtained from *Lithospermum erythrorhizon* in catalyzing the oxidation of geranylhydroquinone during Shikonin biosynthesis. Phytochemistry 175: 112375

Song, W., Zhuang, Y., Liu, T. (2021) CYP82AR Subfamily Proteins Catalyze C-1′ Hydroxylations of Deoxyshikonin in the Biosynthesis of Shikonin and Alkannin. Org. Lett. 23: 2455–2459

Sullivan, M.L. (2015) Beyond brown : polyphenol oxidases as enzymes of plant specialized metabolism. Front. Plant Sci. 5: 783

Suttiyut, T., Auber, R.P., Ghaste, M., Kane, C.N., McAdam, S.A.M., Wisecaver, J.H., et al. (2022) Integrative analysis of the shikonin metabolic network identifies new gene connections and reveals evolutionary insight into shikonin biosynthesis. Hortic. Res. 9: uhab087

Tanaka, S., Tajima, M., Tsukada, M., Tabata, M. (1986) A Comparative Study on Anti-Inflammatory Activities of the Enantiomers, Shikonin and Alkannin. J. Nat. Prod. 49: 466– 469

Tang, C. (2021) Exploring the evolutionary process of alkanin/shikonin *O*-acyltransferases by a reliable *Lithospermum erythrorhizon* genome. DNA Res. 28: 1–9

Tatsumi, K., Yano, M., Kaminade, K., Sugiyama, A., Sato, M., Toyooka, K., et al. (2016) Characterization of shikonin derivative secretion in *Lithospermum erythrorhizon* hairy roots as a model of lipid-soluble metabolite secretion from plants. Front. Plant Sci. 7: 1066

Tatsumi, K., Ichino, T., Onishi, N., Shimomura, K., Yazaki, K. (2020) Highly efficient method of *Lithospermum erythrorhizon* transformation using domestic *Rhizobium rhizogenes* strain A13. Plant Biotechnol. 37: 39–46

Tatsumi, K., Ichino, T., Isaka, N., Sugiyama, A., Moriyoshi, E., Okazaki, Y. (2023) Excretion of triacylglycerol as a matrix lipid facilitating apoplastic accumulation of a lipophilic metabolite shikonin. J. Experomental Bot. 74: 104–117

Tran, L.T., Taylor, J.S., Constabel, C.P. (2012) The polyphenol oxidase gene family in land plants : Lineage-specific duplication and expansion. BMC Genomics 13: 1–12

Ueoka, H., Sasaki, K., Miyawaki, T., Ichino, T., Tatsumi, K., Suzuki, S., et al. (2020) A Cytosol-Localized Geranyl Diphosphate Synthase from *Lithospermum erythrorhizon* and Its Molecular Evolution. Plant Physiol. 182: 1933–1945

Ueta, R., Abe, C., Watanabe, T., Sugano, S.S., Ishihara, R., Ezura, H., et al. (2017) Rapid breeding of parthenocarpic tomato plants using CRISPR/Cas9. Sci Rep 7: 507

Wei, S., Xiang, Y., Zhang, Y., Fu, R. (2022) The unexpected flavone synthase-like activity of polyphenol oxidase in tomato. Food Chem. 377: 131958

Widhalm, J.R., Rhodes, D. (2016) Biosynthesis and molecular actions of specialized 1,4-naphthoquinone natural products produced by horticultural plants. Hortic. Res. 3: 16046

Xu, W., Ma, W., Yue, J., Hu, Y., Zhang, Y., Wang, H., et al. (2025) Phytomedicine Juglone alleviates pelvic pain and prostatic inflammation via inhibiting the activation of NLRP3 inflammasome and alleviating oxidative stress in EAP mice. Phytomedicine 142: 156732

Yamamoto, H., Inoue, K., Li, S., Heide, L. (2000a) P-450 monooxygenase from *Lithospermum erythrorhizon* cell suspension cultures. Planta 210: 312–317

Yamamoto, H., Yazaki, K., Inoue, K. (2000b) Simultaneous analysis of shikimate-derived secondary metabolites in *Lithospermum erythrorhizon* cell suspension cultures by high-performance liquid chromatography. J. Chromatogr. B. Biomed. Sci. Appl. 738: 3–15

Yamazaki, M., Shibata, M., Nishiyama, Y., Springob, K., Kitayama, M., Shimada, N., et al. (2008) Differential gene expression profiles of red and green forms of *Perilla frutescens* leading to comprehensive identification of anthocyanin biosynthetic genes. FEBS J. 275: 3494–3502

Yazaki, K. (2017) *Lithospermum erythrorhizon* cell cultures: Present and future aspects. Plant Biotechnol. 34: 131–142

Yazaki, K., Kunihisa, M., Fujisaki, T., Sato, F. (2002) Geranyl Diphosphate:4-Hydroxybenzoate Geranyltransferase from *Lithospermum erythrorhizon*. CLONING AND CHARACTERIZATION OF A KEY ENZYME IN SHIKONIN BIOSYNTHESIS. J. Biol. Chem. 277: 6240–6246

Yazaki, K., Matsuoka, K., Ujihara, T., Sato, F. (1999) Shikonin Biosynthesis in *Lithospermum erythrorhizon*: Light-induced Negative Regulation of Secondary Metabolism. Plant Biotechnol. 16: 335–342

Yoruk, R., Marshall, M.R. (2003) PHYSICOCHEMICAL PROPERTIES AND FUNCTION OF PLANT POLYPHENOL OXIDASE : A REVIEW. J. Food Biochem. 27: 361–422

Yu, Z., Liao, Y., Zeng, L., Dong, F., Watanabe, N. (2020) Transformation of catechins into thea flavins by upregulation of *CsPPO3* in preharvest tea (*Camellia sinensis*) leaves exposed to shading treatment. Food Res. Int. 129: 108842

Zhang, K., Lu, K., Qu, C., Liang, Y., Wang, R., Chai, Y., Li, J. (2013) Gene Silencing of *BnTT10* Family Genes Causes Retarded Pigmentation and Lignin Reduction in the Seed Coat of *Brassica napus*. PLoS ONE 8: e61247

