## Supplementary Figures and Tables for "CRISPR/Cas9-mediated genome editing reveals the involvement of a polyphenol oxidase in the shikonin-specific biosynthesis in *Lithospermum erythrorhizon*"

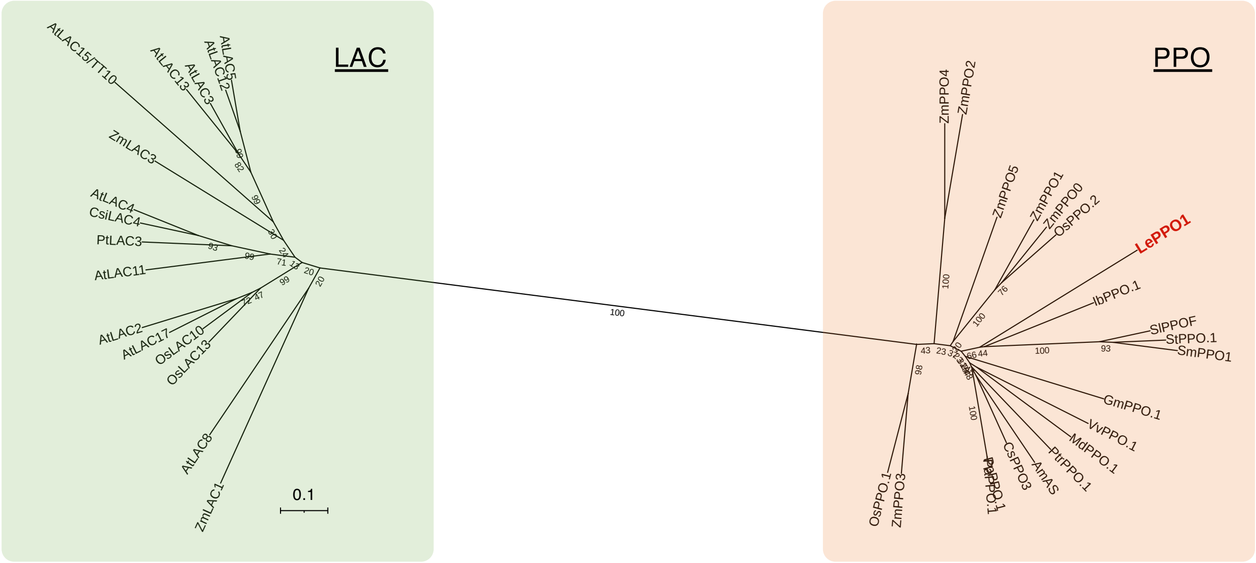


**Supplemental Figure S1.** Phylogenetic relationship of plant polyphenol oxidases (PPOs) and laccases (LACs). The protein sequences were aligned by MUSCLE and Neighbor-joining tree was constructed by MEGA12 (v12.0.10) with bootstrap method (1,000 replications). LePPO1 is represented by red letters. The accession numbers of protein sequences used in this analysis are described in supplementary Table S4. Am, *Antirrhinum majus*; Cs, *Camellia sinensis*; Gm, *Glycine max*; Ib, *Ipomoea batatas*; Le, *Lithospermum erythrorhizon*; Md, *Malus domestica*; Os, *Oryza sativa*; Pa, *Prunus armeniaca*; Pp, *Prunus persica*; Ptr, *Populus trichocarpa*; Sl, *Solanum lycopersicum*; Sm, *Solanum melongena*; St, *Solanum tuberosum*; Vv, *Vitis vinifera*; Zm, *Zea mays*. AS, aureusidin synthase; TT10, transparent testa10.

| | sgRNA4 | sgRNA5 | sgRNA1 | sgRNA3 | ratio |
|---|---|---|---|---|---|
| WT | CCATGTGGTATGACAATTCCAGC | TAACATGGTCTCCAATGCCGTGG | CCATCTTTTATTGCCACCACGGA | GCTAGGTTATGAATACCAGAAGG | 0/10 |
| #112_1 | CCATGTGAGATGG---------- | -------------AATGGGCCGG | ----CGGTTATTGCCACCACGGA | ----------------------- | 7/10 |
| #112_2 | CCATGTG-----------CCAGC | ----------------------- | CCATCTTTTATTGCCACCACGGA | GCTAGGTTATG-----CAGAAGG | 2/10 |
| #112_3 | CCATGTG-----------CCAGC | ----------------------- | CCATCTTTTATTGCCACCACGGA | GCTAGGTTATG-----CAGAAGG | 1/10 |
| WT | CCATGTGGTATGACAATTCCAGC | TAACATGGTCTCCAATG----CCGTGG | CCATCTTTTATTGCCACCACGGA | GCTAGGTTATGAATACCAGAAGG | 0/10 |
| #113_1 | CCATG------------------ | ---------------TG----CCGTGG | CCATCTT---------------- | ----------------CAGAAGG | 7/10 |
| #113_2 | CCATGTGCATTGGAGACCATGTT | -AACATCGCTGGAATTGTCATCCGTGG | CCATCTTTTATTGCCACCACGGA | GCTAGGTTATGAATA-CAGAAGG | 3/10 |
| WT | CCATGTGGTATGACAATTCCAGC | TAACATGGTCTCCAATGCCGTGG | CCATCTTTTATTGCCACCACGGA | GCTAGGTTATGAATACCAGAAGG | 0/10 |
| #117_1 | CCATGTG----------TCCAGC | TAACATGGTCTCCGG-ATCTTGT | CCATCTTTTATTGCCACCACGGA | ----------------------- | 4/10 |
| #117_2 | TGATACCGCTCGCCGCAGCCGAA | TCGTATGTTGTGTGGAATTGTGA | TTCTCTGTACATGGAAATTGGTT | AATCCGCCCAGCAGCGCATTTGG | 6/10 |

**Supplemental Figure S2.** Mutation patterns of *LePPO1* genome-edited lines. The mutations on sgRNAs are shown as red color. WT, wild type.

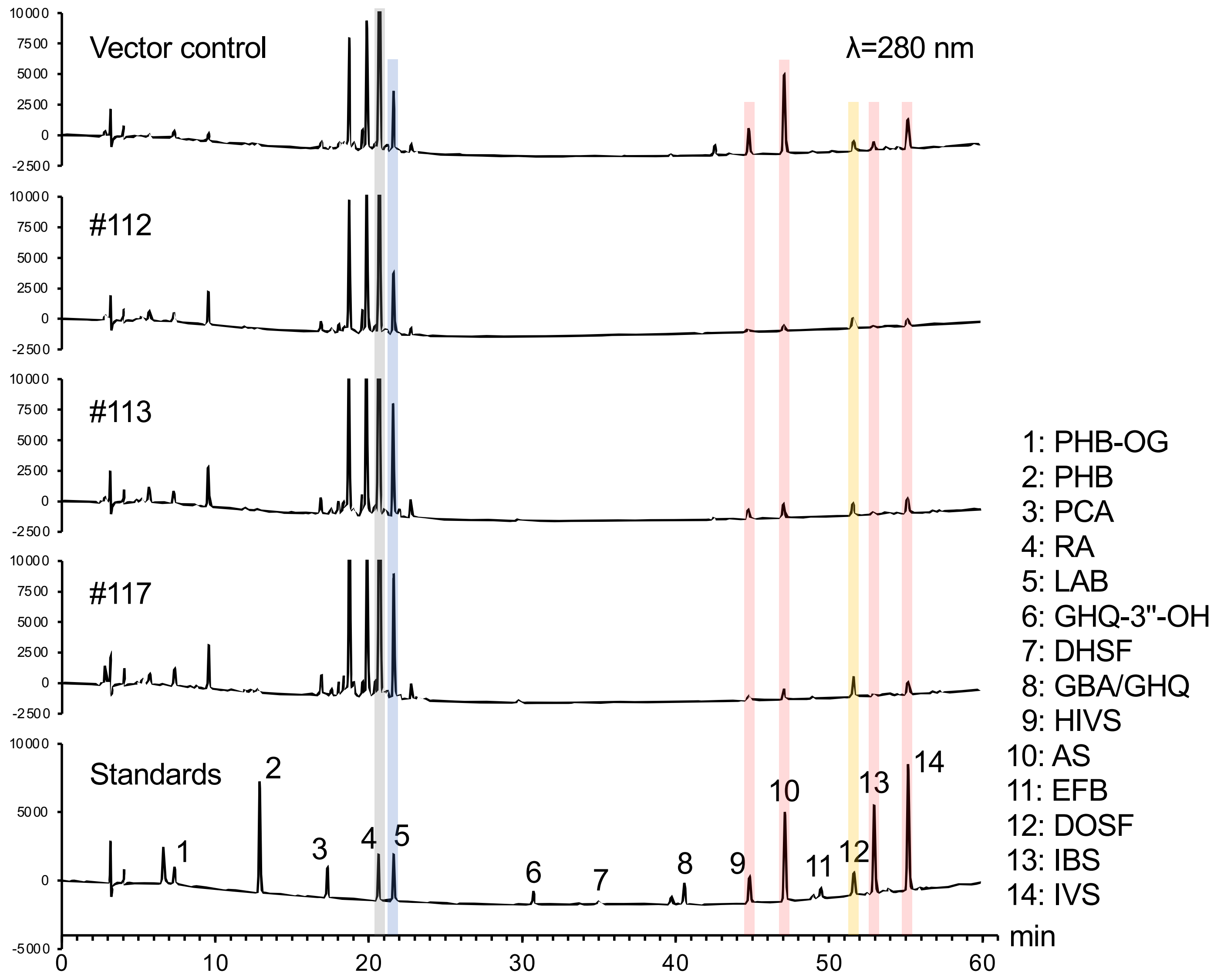


**Supplemental Figure S3.** HPLC chromatograms of *LePPO1* genome-edited lines and vector control (λ=280 nm). PHB-OG, *p*-hydroxybenzoic acid-*O*-glucoside; PCA, *p*-coumaric acid; RA, rosmarinic acid; LAB, lithospermic acid B; GHQ, geranylhydroquinone; DHSF, dihydroshikonofuran; GBA, *m*-geranyl-*p*-hydroxybenzoic acid; HIVS, *β*-hydroxyisovarelylshikonin; AS, acetylshikonin; EFB, echinofuran B; DOSF, deoxyshikonofuran; IBS, isobutyrylshikonin; IVS, isovarelylshikonin.

**Supplementary Table S1.** RNA-seq data used in transcriptome analysis

| Abbreviation in Figure 2 | Sample | SRA number |
|---|---|---|
| HR-L | Hairy root cultures of Lithospermum erythrorhizon under illumination | DRR107693 |
| HR-D | Hairy root cultures of Lithospermum erythrorhizon in the dark | DRR107694 |
| CC-L | Cell cultures of Lithospermum erythrorhizon in M9 medium under illumination | DRR107695 |
| CC-D | Cell cultures of Lithospermum erythrorhizon in M9 medium in the dark | DRR107696 |

**Supplementary Table S2.** Genes used in expression profile analysis

| Gene name | ID in genome data | Accession number in GenBank |
|---|---|---|
| LePAL1 | LE08848.1 | D83075 |
| LeC4H1 | LE02790.1 | AB055507 |
| Le4CL3 | LE08218.1 | LC781614 |
| Le4CL4 | LE04894.1 | LC781615 |
| LeHMGR1 | LE22128.1 | X74783 |
| LeGPPS1 | LE01885.2 | LC427363 |
| LePGT1 | LE18770.1 | AB055078 |
| LePGT2 | LE18614.1 | AB055079 |
| LeGHQH1 (CYP76B100) | LE05331.1 | MN056183 |
| LeGHQH2 (CYP76B101) | LE25459.1 | MN056184 |
| LeDSH1 (CYP82AR) | LE18074.1 | MT921814 |
| LeSAT1 | LE01141.1 | LC520137 |
| LeSAT2 | LE25525.1 | LC747167 |
| LePPO1 | LE31046.1 | This study |
| LePPO2 | LE01843.1 | This study |
| LePPO3 | LE21824.1 | This study |
| LePPO4 | LE16708.1 | This study |
| LePPO5 | LE23862.1 | This study |

**Supplementary Table S3.** Primers used in this study

| Primer name | Sequences (5' to 3') | Description |
|---|---|---|
| LePPO1_3'-RACE | GTAACATGGTCTCCAATGCCG | RACE |
| LePPO1_5'-RACE_1 | CCACGGCATTGGAGACCATGTTACG | RACE |
| LePPO1_5'-RACE_2 | CAGCATCCACGGCATTGGAGACC | RACE |
| LePPO1_full-length_Fw | CCAACATACTGATAGTTGTTATAGTTCAAG | cDNA cloning |
| LePPO1_qPCR_Fw | CAAATGCTGAACAGGTAGACTG | qRT-PCR |
| LePPO1_qPCR_Rv | GGCATTCATGTACTTCCGAAT | qRT-PCR |
| LeACT_qPCR_Fw | CAACTGGGACGACATGGAGA | qRT-PCR |
| LeACT_qPCR_Rv | GAGTCATCTTCTCTCTGTTGGCC | qRT-PCR |
| F4A_tgRNA_LePPO1_gRNA1 | ttgggtctcgTGCAGTCCGTGGTGGCAATAAAAGAGTTTTAGAGCTAGAAATAGCA | genome-editing |
| R4A_tgRNA_LePPO1_gRNA3 | ttgggtctccTCATAACCTAGCTGCACCAGCCGGGAATCGAA | genome-editing |
| F4A_tgRNA_LePPO1_gRNA3 | ttgggtctcgATGAATACCAGAGTTTTAGAGCTAGAAATAGCA | genome-editing |
| R4A_tgRNA_LePPO1_gRNA4 | ttgggtctccGACAATTCCAGCTGCACCAGCCGGGAATCGAA | genome-editing |
| F4A_tgRNA_LePPO1_gRNA4 | ttgggtctcgTGTCATACCACAGTTTTAGAGCTAGAAATAGCA | genome-editing |
| R4A_tgRNA_LePPO1_gRNA5 | ttgggtctccAAACCGGCATTGGAGACCATGTTACTGCACCAGCCGGGAATCGAA | genome-editing |
| LePPO1_genotyping_Fw | GAGGTGAGATTCTCTGTACATGGAAATTGGTTCTTCC | genotyping |
| LePPO1_genotyping_Rv | TCTGCACCAACACCTTGATCACCTTGTCTA | genotyping |

**Supplementary Table S4.** Plant PPOs and LACs used in Supplementary Figure S1

| Protein name | Species | Accession number in GenBank |
|---|---|---|
| AmAS | *Antirrhinum majus* | BAB20048 |
| CsPPO3 | *Camellia sinensis* | QDX19553 |
| GmPPO.1 | *Glycine max* | KRH00631 |
| IbPPO.1 | *Ipomoea batatas* | AAW78869 |
| MdPPO.1 | *Malus domestica* | AFI08581 |
| OsPPO.1 | *Oryza sativa* | BAB89047 |
| OsPPO.2 | *Oryza sativa* | ABG23045 |
| PaPPO.1 | *Prunus armeniaca* | CAB4276004 |
| PpPPO.1 | *Prunus persica* | ONI10336 |
| PtrPPO.1 | *Populus trichocarpa* | AAU12256 |
| SlPPO_F | *Solanum lycopersicum* | CAA78300 |
| SmPPO1 | *Solanum melongena* | ACT22523 |
| StPPO.1 | *Solanum tuberosum* | AAA85121 |
| VvPPO.1 | *Vitis vinifera* | AAB41022 |
| ZmPPO_0 | *Zea mays* | PWZ46049 |
| ZmPPO_1 | *Zea mays* | PWZ45673 |
| ZmPPO_2 | *Zea mays* | PWZ26247 |
| ZmPPO_3 | *Zea mays* | PWZ06435 |
| ZmPPO_4 | *Zea mays* | PWZ43422 |
| ZmPPO_5 | *Zea mays* | PWZ33588 |
| AtLAC2 | *Arabidopsis thaliana* | NP_180477 |
| AtLAC3 | *Arabidopsis thaliana* | NP_180580 |
| AtLAC4 | *Arabidopsis thaliana* | NP_565881 |
| AtLAC5 | *Arabidopsis thaliana* | NP_181568 |
| AtLAC8 | *Arabidopsis thaliana* | NP_195724 |
| AtLAC11 | *Arabidopsis thaliana* | NP_195946 |
| AtLAC12 | *Arabidopsis thaliana* | NP_196158 |
| AtLAC13 | *Arabidopsis thaliana* | NP_196330 |
| AtLAC15/TT10 | *Arabidopsis thaliana* | NP_199621 |
| AtLAC17 | *Arabidopsis thaliana* | NP_200810 |
| CsiLAC4 | *Citrus sinensis* | XP_006480970 |
| OsLAC10 | *Oryza sativa* | NP_001404427 |
| OsLAC13 | *Oryza sativa* | P0DKK6 |
| PtrLAC3 | *Populus trichocarpa* | CAC14719 |
| ZmLAC1 | *Zea mays* | AAX83112 |
| ZmLAC3 | *Zea mays* | CAJ30499 |

**Supplementary Table S5.** Plant PPOs used in this study

| Protein name | Species | Accession number in GenBank |
|---|---|---|
| CsPPO1 | *Camellia sinensis* | QDX19551 |
| CsPPO2 | | QDX19552 |
| CsPPO3 | | QDX19553 |
| GmPPO.1 | *Glycine max* | KRH00631 |
| GmPPO.2 | | KRH00632 |
| GmPPO.3 | | KRH10820 |
| GmPPO.4 | | KRH10822 |
| GmPPO.5 | | KRH20505 |
| GmPPO.6 | | KRH20508 |
| GmPPO.7 | | KRH21500 |
| GmPPO.8 | | KRH21501 |
| GmPPO.9 | | KRH55663 |
| GmPPO.10 | | KRH62648 |
| IbPPO.1 | *Ipomoea batatas* | AAW78869 |
| IbPPO.2 | | CAC29040 |
| MdPPO.1 | *Malus domestica* | AFI08581 |
| MdPPO.2 | | AFI08582 |
| MdPPO.3 | | AFI08583 |
| MdPPO.4 | | AFI08584 |
| MdPPO.5 | | BAA21676 |
| MdPPO.6 | | BAA21677 |
| PaPPO.1 | *Prunus armeniaca* | CAB4276004 |
| PaPPO.2 | | CAB4276009 |
| PaPPO.3 | | CAB4276010 |
| PaPPO.4 | | CAB4276011 |
| PaPPO.5 | | CAB4276012 |
| PaPPO.6 | | CAB4276013 |
| PaPPO.7 | | CAB4290256 |
| PpPPO.1 | *Prunus persica* | ONI10336 |
| PpPPO.2 | | ONI10337 |
| PpPPO.3 | | ONI10339 |
| PpPPO.4 | | ONI10340 |
| PpPPO.5 | | ONI10341 |
| PtrPPO.1 | *Populus trichocarpa* | AAU12256 |
| PtrPPO.2 | | AAG21983 |
| PtrPPO.3 | | AEH41424 |
| PtrPPO.4 | | AEH41425 |
| PtrPPO.5 | | PNT59063 |
| PtrPPO.6 | | PNT59064 |
| SlPPO A | *Solanum lycopersicum* | CAA78295 |
| SlPPO B | | CAA78296 |
| SlPPO C | | CAA78297 |
| SlPPO D | | CAA78298 |
| SlPPO E | | CAA78299 |
| SlPPO F | | CAA78300 |
| SmPPO1 | *Solanum melongena* | ACT22523 |
| SmPPO2 | | ADG56700 |
| SmPPO3 | | ADY18409 |
| SmPPO4 | | ADY18410 |
| SmPPO5 | | ADY18411 |
| SmPPO6 | | ADY18412 |
| StPPO.1 | *Solanum tuberosum* | AAA85121 |
| StPPO.2 | | AAA85122 |
| VvPPO.1 | *Vitis vinifera* | AAB41022 |
| VvPPO.2 | | CAN61652 |
| VvPPO.3 | | CAN62983 |
| AmAS | *Antirrhinum majus* | BAB20048 |
| AtLAC15 (TT10) | *Arabidopsis thaliana* | NP_199621 |
